# Wax worm saliva and the enzymes therein are the key to polyethylene degradation by *Galleria mellonella*

**DOI:** 10.1101/2022.04.08.487620

**Authors:** A. Sanluis-Verdes, P. Colomer-Vidal, F. Rodríguez-Ventura, M. Bello-Villarino, M. Spinola-Amilibia, E. Ruiz-López, R. Illanes-Vicioso, P. Castroviejo, R. Aiese Cigliano, M. Montoya, P. Falabella, C. Pesquera, L. González-Legarreta, E. Arias-Palomo, M. Solà, T. Torroba, C.F. Arias, F. Bertocchini

**Author notes:** Equal contribution (A. Sanluis-Verdes, P. Colomer-Vidal). Corresponding authors: CFA, FB.

## Abstract

Plastic degradation by biological systems with re-utilization of the by-products can be the future solution to the global threat of plastic waste accumulation. We report that the saliva of *Galleria mellonella* larvae (wax worms) is capable of oxidizing and depolymerizing polyethylene (PE), one of the most produced and sturdy polyolefin-derived plastics. This effect is achieved after a few hours’ exposure at room temperature and physiological conditions (neutral pH). The wax worm saliva can indeed overcome the bottleneck step in PE biodegradation, that is the initial oxidation step. Within the saliva, we identified two enzymes that can reproduce the same effect. This is the first report of enzymes with this capability, opening up the way to new ground-breaking solutions for plastic waste management through bio-recycling/up-cycling.

## Introduction

Polyethylene (PE) accounts for 30% of synthetic plastic production, largely contributing to plastic waste pollution on the planet to-date [1]. Together with polypropylene (PP), polystyrene (PS) and polyvinylchloride (PVC), PE is one of the most resistant polymers, with very long C-C chains organized in a crystalline, dense structure. Given the hundreds of million tons of plastic waste accumulating and the still escalating pace of plastic production, re-utilization of plastic residues is a necessary path to alleviate the gravity of the plastic pollution problem, and at the same time to render available a huge potential reservoir of carbon [2]. To-date, only mechanical recycling is being applied at a large scale. Several factors, such as the low number of plastic types prone to be mechanically recycled, and the low quality of the secondary products severely restrict the potential of this solution to the problem of plastic waste accumulation. Chemical recycling, as an alternative procedure, is preferentially aiming at plastic upcycling, e.g. decomposing polyolefin-derived plastics in order to take advantage of smaller intermediates. Several technologies have been applied at a lab scale, although the high energetic cost might still impede the scaling up of these technological tools [3].

In addition to mechanical and chemical recycling, biodegradation is widely considered as a promising strategy to dispose of plastic residues. Biodegradation refers to environmental degradation by biological agents. As indicated by the IUPAC definition [4] it is defined as the “breakdown of a substance catalyzed by enzymes in vitro or in vivo”, later modified “to exclude abiotic enzymatic processes” [5]. In the case of PE, biodegradation requires the introduction of oxygen into the polymeric chain [6, 7]; this causes the formation of carbonyl groups and the subsequent scission of the long hydrocarbon chains with production of smaller molecules, which can then be metabolized by microorganisms [8, 9]. The crucial first step of this chain of events, i.e. the oxidation of the PE polymer, is usually carried out by abiotic factors such as light or temperature [6, 8, 10]. Once the long polymeric molecules are broken down, a process that takes years of exposure to environmental factors in the wild, bacteria or fungi intervene and continue the job [6, 9, 11, 12]. This is the current paradigm driving the research field in biodegradation. Within this paradigm, several bacterial and fungal strains have been identified as capable to carry on a certain extent of PE degradation. However, in most of the cases such degradation requires an aggressive pre-treatment of PE (heating, UV light, etc.) that accelerates the incorporation of oxygen into the polymer, making the abiotic oxidation the real bottleneck of the reaction [12–16]. In the past decade a few microorganisms have been described as capable to act on untreated PE [17–23], although they require significantly longer incubation times compared to experimental conditions with pre-oxidized PE.

The identification of enzymes from microorganisms capable of degrading untreated PE has proven a much difficult task. In fact, no such enzyme has been identified yet, confirming the crucial limiting role of oxidation in the whole biodegradation process chain [12]. Reported enzymes capable of acting on polyolefin-derives plastics require a pretreatment of the plastic material [12, 14]. For example, two reported laccases, able to chemically modify PE, necessitate an abiotic pretreatment [24] or the addition of redox mediators such as 1-hydroxybenzotriazole [25].

This scenario confirms that the synthetic nature of the compound, together with the hydrophobicity and inaccessibility features, make plastic a difficult target for animal, fungal or microbial-derived enzymatic activities. Nonetheless, some lepidopteran and coleopteran insects revealed the unexpected capacity to degrade untreated PE and PS [26–31]. The larvae of *Galleria mellonella*, also known as wax worms (ww), can oxidize PE within one hour from exposure [29, 32–38], making it the fastest known biological agent capable of chemically modifying PE.

From the original observation that plastic debris appears when PE film is in contact with the recently formed ww cocoon and mouth secretion, we analyzed the saliva of the ww (GmSal), which revealed its capacity to oxidize and break PE within a time frame of a few hours. This effect is confirmed by the GPC analysis of GmSal-treated PE, showing the scission of long hydrocarbon chains into small molecules. Using Gas Chromatography-Mass Spectrometry (GC-MS) we identified degradation products such as small oxidized aliphatic chains, further confirming the breaking of the aliphatic chain polymer in shorter molecules. Proteomic analyses of GmSal revealed the presence of a handful of enzymes belonging to the hexamerin/prophenoloxidase family. Two of these, an arylphorin re-named Demetra, and an hexamerin re-named Ceres oxidize PE after a few hours’ application at room temperature (RT), an effect accompanied in the case of Demetra by the physical deterioration of PE and the release of degradation by-products similar to the ones obtained with the whole saliva.

Demetra and Ceres are the first enzymes capable of producing such modifications on a PE film working at room temperature and in a very short time, embodying a promising alternative to the abiotic oxidation of plastic, the first and most difficult step in the degradation process. The identification of invertebrate enzymes capable of oxidizing PE in a few hours represents a totally new paradigm in the world of plastic degradation and more widely in the plastic waste management fields, and opens up a highway of possibilities to solve the plastic waste pollution issue, together with the designing of new formulae/routes for synthetic polymers production.

## Results

### Wax worm saliva oxidizes PE film

Saliva, broadly defined here as the juice present in the anterior portion of the digestive apparatus, was collected from the ww mouth and tested on a commercial PE film (Fig. 1A). After three consecutive applications of 30 *μ*l of GmSal for 90 minutes each, Confocal Raman microscopy/Raman spectroscopy (RAMAN) analysis indicated polymer oxidation, accompanied by a general deterioration of the film (Fig. 1B). This is evident in the overlapping with the PE control (Figs. 1C, D), which reveals the expected PE signature profile. As a further control, the saliva of another lepidopteran larva, *Samia cynthia*, was applied on the PE film, and no oxidation was generated (Figure 1E). The changes produced by the GmSal in a few hours-long applications are similar to those generated by environmental factors after months or years of exposure to weathering [39, 40]). The Fourier Transformed Infrared Spectroscopy (FTIR) analysis confirmed the oxidation profile (Fig S1). The changes in PE chemical composition revealed by the spectroscopy techniques suggested that molecules other than the long PE polymeric chain formed upon the contact with GmSal.

**Figure 1.**
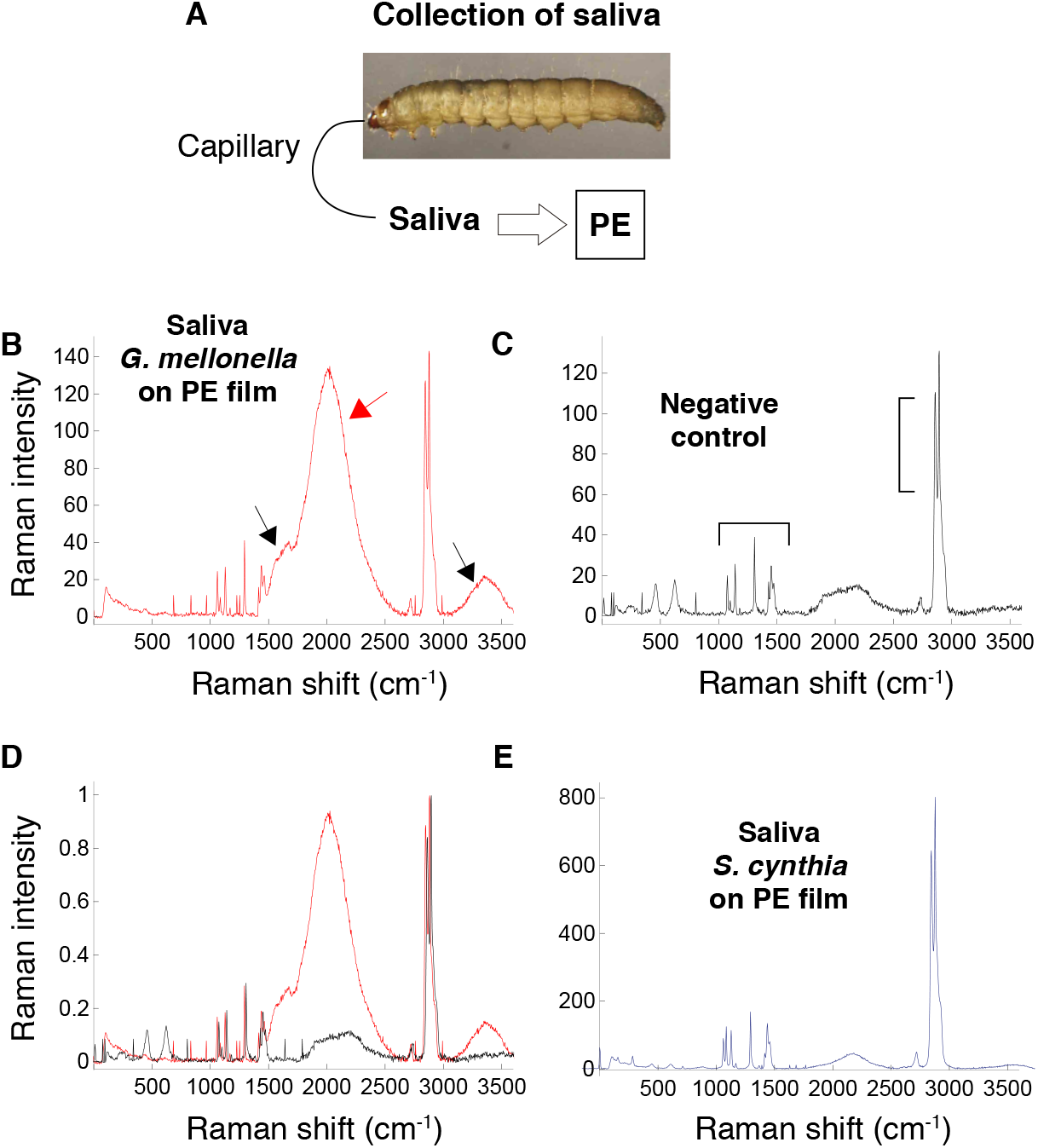
*Galleria mellonella* saliva (GmSal) collection and functional study. A. Scheme of saliva collection and application. B, C, D, E. RAMAN analysis of PE film. B. PE film treated with GmSal: three applications of 90 minutes, 30 *μ*l each. The peaks between 1500 and 2400 cm^−1^ indicate different collective stretching vibrations due to the presence of other organic compounds, sign of PE deterioration (red arrow). Oxidation is indicated between 1600 and 1800 cm^−^1 (carbonyl group) and 3000-3500 cm^−^1 (hydroxyl group) (black arrows) [41]). C. Control PE film. Brackets indicate the picks that characterize PE (PE signature), corresponding to the bands at 1061, 1128, 1294, 1440, 2846 and 2880 cm^−^1. D. Overlapping profiles (B and C). E. PE film treated with *Samia cynthia* saliva.

### Wax worm saliva degrades PE

The molecular weight characteristics of PE before and after treatment with GmSal were analysed using High Temperature-Gel Permeation Chromatography (HT-GPC). After a few hours’ applications of GmSal on PE film (15 applications, 90 minutes each), the molecular weight distribution became bi-modal, with a new peak in the low molecular weight region, indicating the breaking of the C-C bonds with the appearance of small compounds (Figure 2A). Formation of C=O bonds as a consequence of chain scission was shown by the increase of the carbonyl index (CI, Figure 2A). The weight average (*M*_*w*_) was only slightly changed (from 207,100 to 199,500 g/mol) suggesting that the polymer was not uniformly modified by the GmSal, with portions that have been strongly depolymerized, while others remained still untouched [42]. The same result was obtained using PE 4000 instead of PE film, with a change in *M*_*w*_ from 4,000 to 3,900 g/mol and an increase in the carbonyl index in the GmSal-treated sample (Figure 2B, black and blue curves).

**Figure 2.**
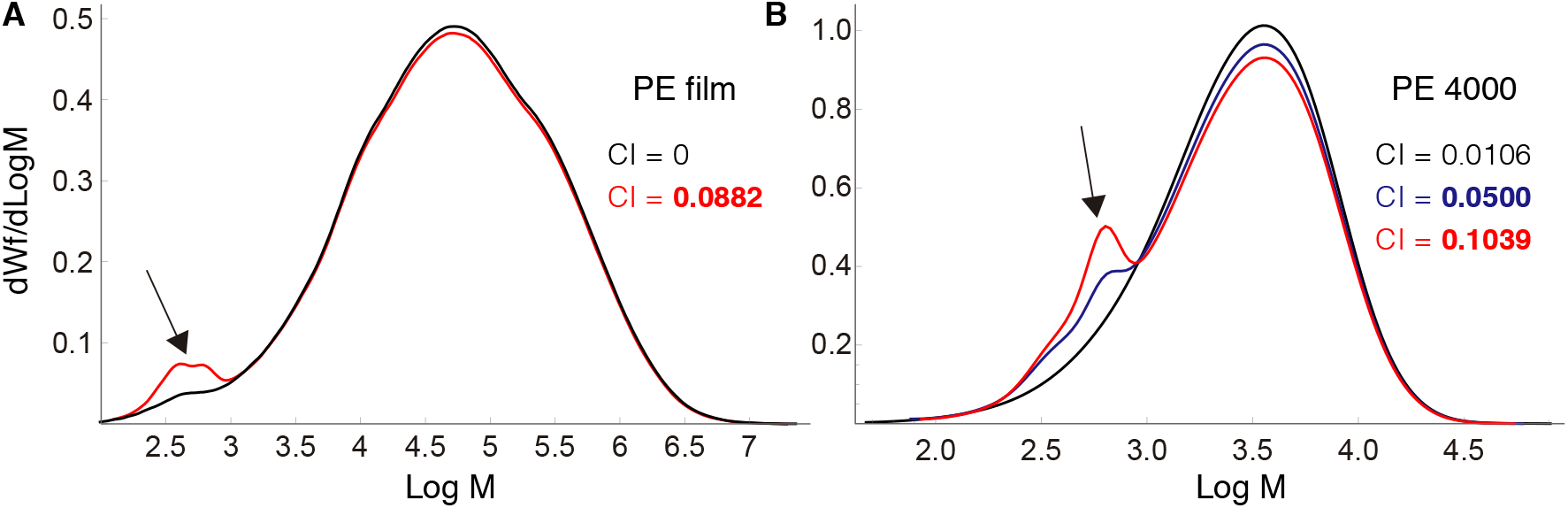
HT-GPC analyses of PE treated with GmSal. Molecular weight distribution of PE film (A) and PE 4000 (B) are indicated. A. Control PE (in black) and PE-treated (in red) are compared. The carbonyl index (CI) of the control and experimental are indicated. B. Control PE (black), treated-PE (15 applications) (blue), treated-PE (30 applications) are compared. The CI of each sample is indicated. Arrows indicate compounds with low molecular weight in the treated PE.

These results show that PE was oxidized and depolymerized as a consequence of GmSal exposure, with formation of oxidized molecules of low molecular weight. In order to analyse the dynamics of changes in the PE after increased exposure to GmSal in time, we doubled the exposure time of PE 4000 to GmSal (30 applications, 90 minutes each) (Figure 2B, red curve). Breaking of large polymer chains with formation of smaller molecules notably increased at longer exposure times, as showed by the comparison of the molecular weight distributions and by the doubling of the CI (Figure 2B). These experiments confirmed the PE oxidation with depolymerization upon a few hours’ exposure to GmSal, an effect that increased with time, and pointed to the presence therein of still unknown activities capable of PE degradation.

### Identification and analysis of the by-products of PE treated with wax worm saliva

To have an insight into the degradation products resulting from PE-saliva contact and released from the oxidized polymer, PE granules (crystal polyethylene-PE 4000) were exposed to GmSal and subsequently analysed by Gas Chromatography-Mass Spectrometry (GC-MS) and identified by NIST11 library for untargeted compounds. After 9 applications of 40 *μ*L of GmSal for 90 minutes each at RT, new compounds were detected in the experimental sample (Fig. 3).

**Figure 3.**
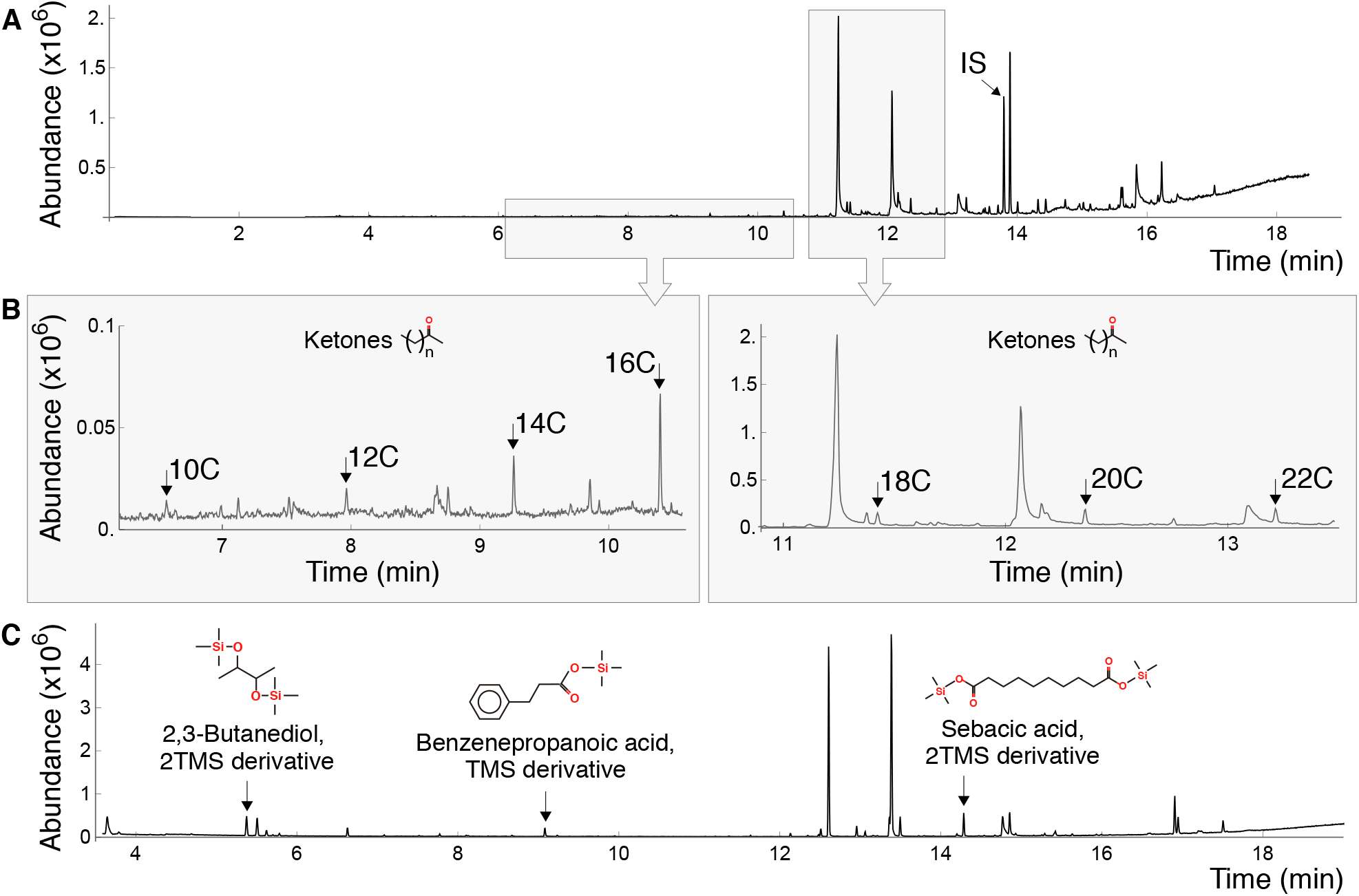
Identification of PE degradation by-products via GC-MS. A, B, C. Chromatograms of PE treated with GmSal, indicating different compounds. A and B. Ketones of different length, indicated by the number of carbon atoms. C. 2,3-Butanediol 2TMS derivative, benzenepropanoic acid TMS derivative, sebacic acid sTMS derivative. Compounds in C were identified with silylation (see Methods). IS: internal standard.

The detected compounds comprised oxidized aliphatic chains, like 2-ketones from 10 to 22 carbons. Ketones from 10 to 18 carbons were identified by comparing the fragmentgram of the ion m/z 58 from methyl ketones that correspond with the transposition of the McLafferty on the carbonyl located at the second carbon of each ketone. Those not present in the library as 2-eicosanone and 2-docosanone showed the same fragmentgram m/z 58 and were defined by the equidistance of the peaks along the retention time and their molecular weight (Table S1). Furthermore, the presence of 2-ketones was confirmed with GC-MS/MS in MRM mode with the ion with the highest m/z and the exclusive of each molecule as 212 > 58 for 2-tetradecanone, 240 > 59 for 2-hexadecanone, and 282 > 58 for 2-octadecanone. Also, butane, 2,3-Butanediol, 2-trimethylslyl (TMS) derivative, and sebacic acid, 2TMS derivative were identified using sample silylation, indicating the deterioration of the PE chain (Fig 3C). At the same time, a small aromatic compound recognizable as benzenepropanoic acid, TMS derivative, a plastic antioxidant, was found. Derivative chemicals were confirmed as well using GC-MS/MS with an m/z of 147 > 73, 331 > 73, and 104 > 75, respectively.

The presence of this plastic antioxidant suggests an “opening” in the polymeric structures, with the release of small stabilizing compounds normally present in plastics (plastic additives). To verify if an increase in time exposure to GmSal caused an increase in PE degradation, we repeated the experiment of PE 4000 exposure in sequential times, with four applications per day of 100 *μ*L of GmSal for 90 minutes each at RT, performed in 1, 2, 3 and 6 consecutive days. The analysis of the supernatants revealed a progressive increase in the formation of 2-decanone, 2-dodecanone, 2-tetradecanone, and 2-hexadecanone (Fig S2). These data indicate an increase of at least twice the relative abundance of degradation products with time (from 1 to 6 days) as a consequence of prolonged exposure to the GmSal.

### Study of the wax worn saliva: enzymes identification and functional studies

To understand the nature of the buccal juice, a GmSal sample was analysed by negative staining electron microscopy (EM), revealing a high content of proteins or protein complexes (size:10-15 nm) (Fig. 4A). No other structures (such as vesicles or bacteria) were detected. Electrophoretic analysis (SDS-PAGE) confirmed the presence of proteins with a prominent band at around 75kDa (Fig. 4B).

**Figure 4.**
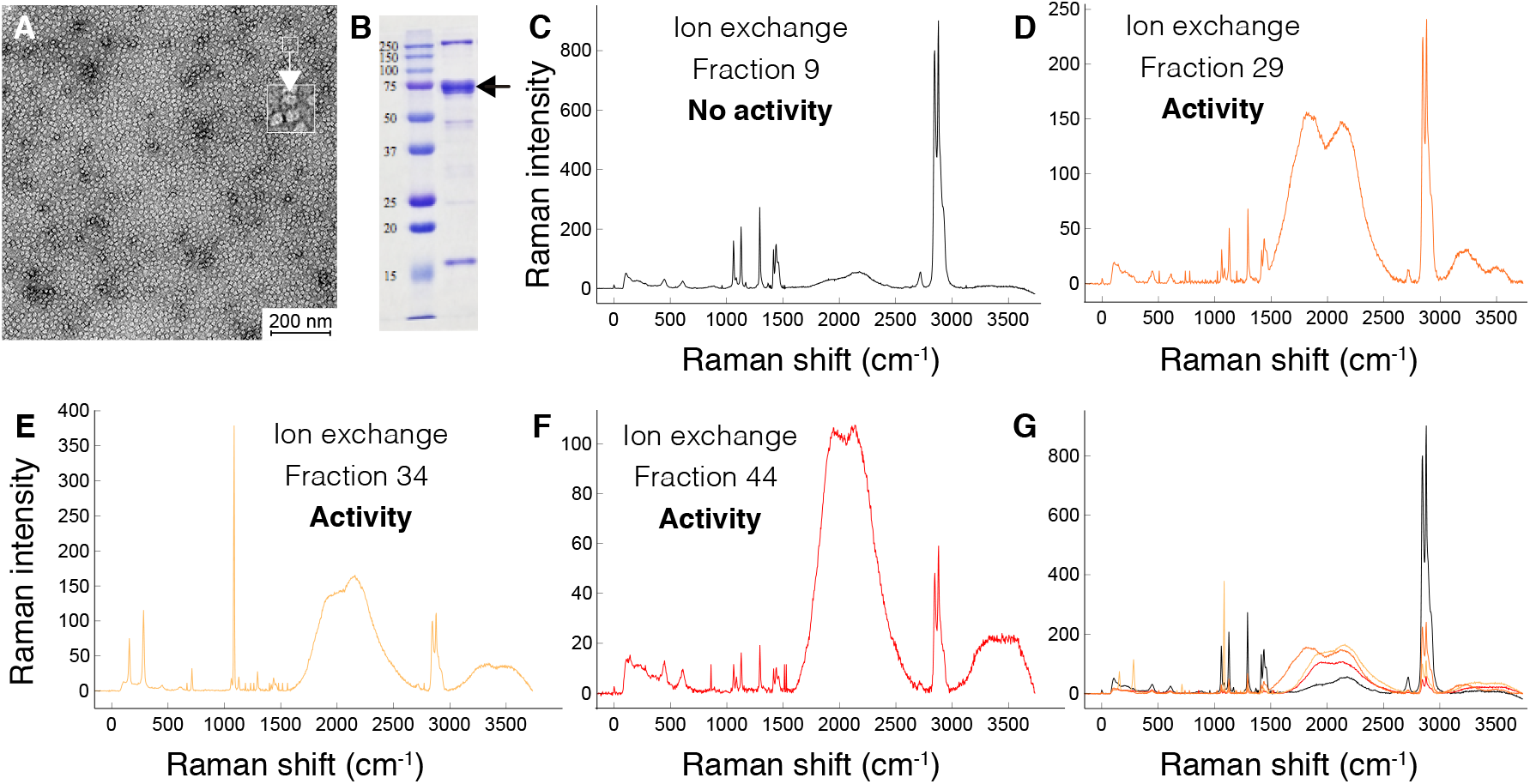
GmSal content functional characterization. A. Electron microscopy negative staining of a saliva sample, dilution 1:500. Protein complexes (10-15 nm size) are indicated in the top right square. B. SDS-PAGE of a GmSal sample, dilution 1:50. Molecular weight standards are listed on the left. The arrow indicates the major band at 75kDa. C-F. RAMAN of PE treated with IEX fractions (see Fig. S3). C. Fraction 9, no activity (control PE film, see Fig. 1 for details). D-F. Profile corresponding to IEX peak 1 (fraction 29), 2 (fraction 34) and 3 (fraction 44) (see Fig. 1 for details). The intense peak at 1085 cm^−^1 in E indicates amorphous PE. The decrease of the peaks at 2845 and 2878 cm^−^1 indicates a less significant presence of PE. G. Overlapping of fractions C to F.

Does GmSal contain enzymes responsible for the detected PE modifications? To assess this, a proteomic analysis of GmSal contents was carried out. More than 200 proteins were detected, including a variety of enzymatic activities, transport and structural proteins, etc. (not shown). To narrow down the number of potential candidates, a saliva sample was analyzed by size exclusion chromatography (SEC). The elution profile showed a main single, wide peak (Fig. S3A). SDS-PAGE gel of the major fraction showed a strong band at about 75kDa (Fig. S3C). The proteomic profile of this band revealed the presence of proteins known in arthropods as related to transport or storage (not shown).

To further identify if the wide peak contained subspecies, an ion exchange chromatography (IEX) was run with a second saliva sample, which showed four well-defined elution peaks (peaks 1 to 4 in Fig. S3B). Analysis by SDS-PAGE indicated that they all contained proteins of similar molecular weight (Fig. S3D). To check which protein fractions of both the SEC and IEX retained PE degradation activity, aliquots of the eluted fractions were tested on a PE film. Using RAMAN spectroscopy, degradation activity was analysed from fractions of the IEX four peaks (Fig. 4C-G) and in the SEC major peak (Fig S4). Peaks indicated as 1, 2 and 3 (as in Fig. S3B) showed a major extent of PE oxidation than fraction 4, which evidently contained some extent of residual activity (not shown). High degradation activity was also detected in the SEC main peak (peak 5 in Fig. S3A) (Fig S4).

Proteomics of IEX peaks 1, 2, and 3 revealed the presence of a handful of proteins, belonging to the arthropodan hexamerin/prophenoloxidase superfamily (peaks 1, 2, and 3, respectively, respectively, proteins’ list not shown). This result on the one hand confirmed the outcome of the SEC fraction proteomics (peak 5), and on the other refined it, reducing the number of potential candidates present in each peak. The fact that this family comprehends oxidase activities, made them the obvious candidates for PE degradation capacity within GmSal. These proteins, namely arylphorin subunit alpha, arylphorin subunit alpha-like, and the hexamerin acidic juvenile hormone-suppressible protein 1 were produced using a recombinant expression system and tested for this ability.

### Identification of wax worm enzymes as PE oxidizers

In order to assess the activity of these proteins, 5 *μ*l of each purified enzyme at a concentration of 1-5 *μ*g/*μ*l, were applied separately 8 sequential times (90 minutes each) on PE films. While arylphorin subunit alpha did not show any effect on PE film (not shown), arylphorin subunit alpha-like, re-named Demetra (NCBI accession number: XP 026756396.1), caused PE deterioration with occasional conspicuous visual effect on the film itself (Fig. 5A-E and Fig. S5). RAMAN confirmed oxidation and a damaged PE signature (Fig. 5F-H). The hexamerin, re-named Ceres (NCBI accession number: XP 026756459.1), caused PE oxidation/deterioration (Fig. S6), but without the visual effect caused by Demetra. The same experiment performed with inactivated enzymes did not show any modification on the PE film, as indicated by RAMAN analysis (Fig. S7). These results support the original hypothesis based on the idea that the buccal secretion is the main source of plastic degradation activities in the ww.

**Figure 5.**
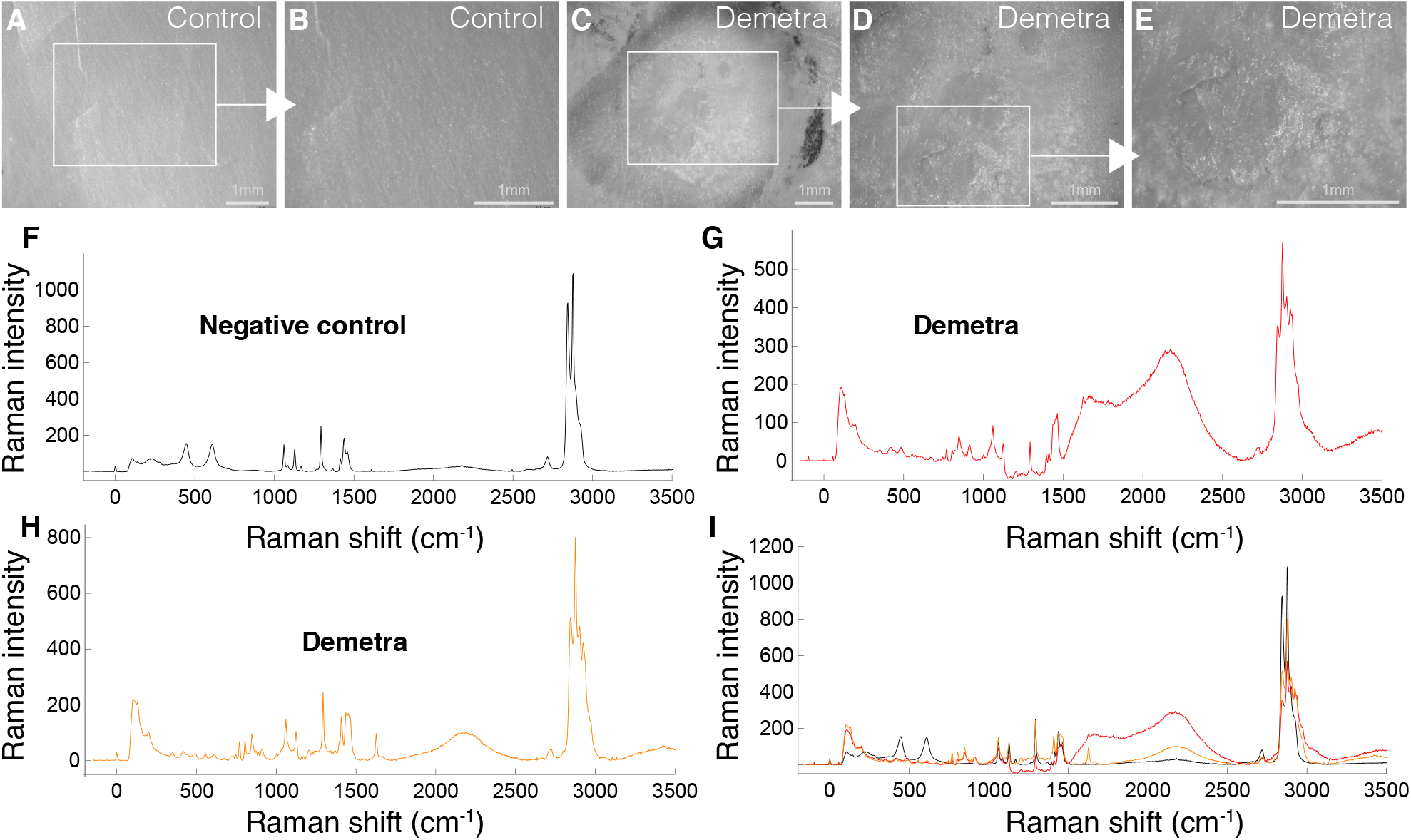
Demetra effect on PE film. A, B. Control PE film. C-E. Demetra on PE film. F-I. RAMAN spectroscopy of control (F) and Demetra treated PE film (G, H). F. Control PE film (see Fig. 1 for details). G, H. Two different punctual analyses within the crater as showed in E, indicating PE deterioration. See Fig. 1 for details. Moreover, the bands at 845 and 916 cm^−^1 correspond to C-O-C and C-COO groups, respectively. The presence of PE is insignificant in G.

To analyse the potentiality of the saliva proteins in oxidizing PE, GC-MS was performed on PE granules (PE 4000) exposed to Demetra or Ceres. After 24 applications of Demetra (10 *μ*l at 1.2 mg/mL, 90 minutes each), 2-ketones from 10 to 22 carbons were detected in the supernatant using GC-MS, the fragmentgram m/z 58, and retention time for identifications (Fig. 6A). Increasing the treatment (ten versus five applications, 90 minutes each) showed an increase of 2-ketones of 10 to 20 carbons in relative abundance, and the appearance of 2-docosanone which was not detected after five applications (Fig. 6B). On the other hand, after PE 4000 treatment with Ceres, no by products were detected in the supernatant, suggesting substantial difference between the two proteins, despite they both shared the capacity to oxidize PE.

**Figure 6.**
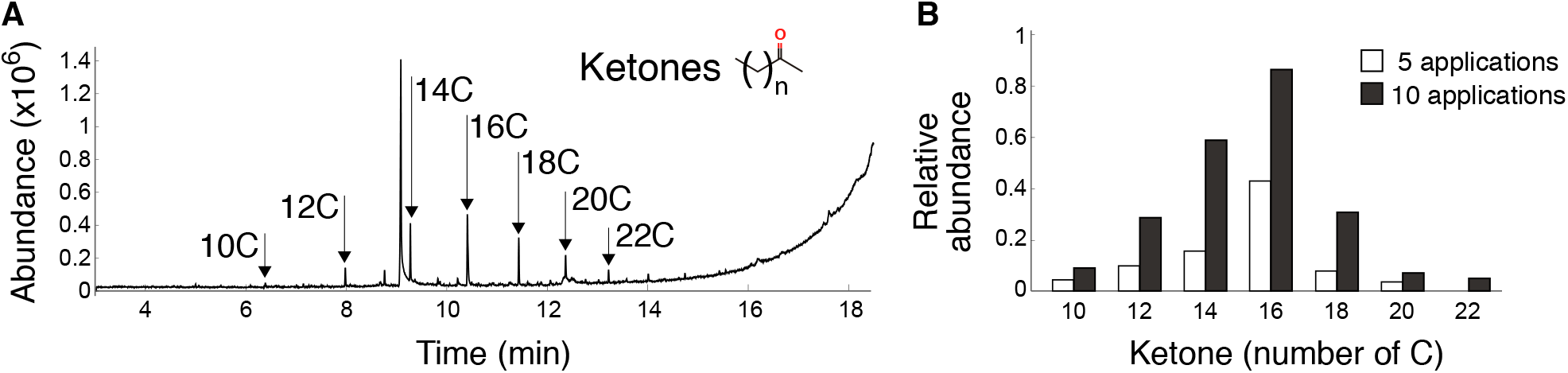
Generation of PE degradation by-products by Demetra. A. GC-MS chromatogram of PE treated with Demetra. The arrows indicated the peaks corresponding to ketones with different number of carbons. B. Increase of ketones formation as degradation products from five to ten applications of Demetra to PE.

## Discussion

This study reports that the saliva of the ww oxidizes and depolymerizes PE, with ww enzymes therein capable of reproducing the effect observed with the whole saliva. This is the first report of an enzymatic activity capable of attacking the PE polymer without any previous abiotic treatment. This capacity is achieved by animal enzymes working at room temperature and in aqueous solution with a neutral pH. Under these conditions, the enzymatic action of the ww saliva overcomes in a few hours a recognized bottleneck step (i.e. oxidation) in PE degradation [6, 8–10].

The capacity of *Galleria mellonella* as well as other Coleoptera and Lepidoptera to degrade sturdy polyolefin-derived polymers as PE or PS has been extensively documented in the past few years, with no pre-treatment required for the plastic polymer to be degraded [26–29, 31–38, 43–48]. If this capacity resides in the microorganisms of the worm gut, in the invertebrate itself, or in a complementation of the two, is still an object of debate. The gut microbiome has traditionally been considered the culprit of plastic degradation by insects. However, despite the numerous reports appeared lately about microbe species being potentially responsible for insect-driven plastic biodegradation, no consensus on specific species or genera of bacteria/fungi colonizing the Lepidoptera and Coleoptera gut and involved in plastic degradation has been reached [26–28, 30, 31, 34–38, 44]. Indeed, the exclusive involvement of microorganisms in this process has been recently questioned [32, 49]. As for *Galleria*, the fast degradation makes it improbable for the gut microbiota to be the sole player in the observed modification of PE chemical structure, as already suggested [32]. This study provides solid evidence in favor of this argument and confirms the key role played by the larvae of *G. mellonella* in PE degradation: the saliva of the ww, and the enzymes it contains, oxidizes and depolymerizes PE. The action of the enzymes present in the saliva of *G. mellonella* on PE is therefore equivalent to that of abiotic pretreatments. The saliva-dependent oxidation of PE could thus provide a suitable substrate for further biological attack by causing the scission of the long polymer chains into smaller molecules that could then be metabolized and assimilated along the insect’s digestive system (by the microbiome and/or by the insect’s cells).

The current paradigm/hypothesis of PE biodegradation stands on the breaking of the C-C bond via the same mechanism that bacteria deploy to break alkanes [50, 51]. However, the two enzymes of the ww saliva identified in this study suggest the potential existence of alternative mechanisms. These enzymes, Demetra and Ceres, are an arylphorin and an hexamerin, respectively [52]. They are phylogenetically related to phenol-oxidases (enzymes targeting aromatic rings) and hemocyanins, oxygen transport proteins that also present phenoloxidase activity [53, 54]. The presence of this type of enzyme in the buccal cavity of insects is not a novelty: insect saliva is a source of enzymes involved in digestion, defence, transport etc. [55, 56]. For example, leaf-feeding lepidopteran larvae secrete saliva on the leaves before ingestion [55, 57]. In this context, the presence of phenol-oxidase-like activities is not surprising, considering that phenols are widely employed by plants as a defence against potential enemies [58–60].

Unexpected instead is the capacity of these particular proteins to oxidize PE, a polymeric, compact hydrophobic substance. However, the ecological niche of Lepidoptera and the potential necessity to react to plant phenols might provide a possible explanation. The appearance of a small aromatic compounds recognizable as a plastic additive such as benzenepropanoic acid, TMS derivative among the degradation products raises the possibility that this compound could become the target of the phenoloxidase-like activity of the ww enzymes. A consequence of this potential action of the ww enzymes on aromatic additives could be the formation of free radicals leading to the initiation of the autoxidative chain reaction [6]. The idea of the formation of free radicals as a first step to trigger autooxidation is not new, having been proposed as a way in which some bacteria might initiate plastic biodeterioration, via still unknown enzymes [14, 20]. However, this will remain a speculation until the presence/formation of radical is verified. Alternative scenarios include a direct enzymatic action on the hydrocarbon chain in some still unknown fashion, or the classical *β* -oxidation mechanism aforementioned. Further studies will be required to get deeper insights into the functional modality of these ww enzymes, their diversity (as already indicated by the differences in the experimental outcomes between Demetra and Ceres), and the molecular mechanisms acting on PE.

The existence of enzymes produced by insects, secreted from the mouth and evolved to work at room temperature and neutral pH on plastic provides a new paradigm for biological degradation of PE. This new framework goes well beyond the current definition of biodegradation, which is exclusively based on the full conversion of plastic to CO_2_ through the metabolic activity of microorganisms: on one hand, the observed oxidation and deterioration of PE do not depend on any microbial activity; on the other hand, the easy working conditions and the appearance of degradation products such as ketones and additives suggest the potential use of these enzymes for plastic waste degradation and recycling or upcycling of plastic components. This potentiality could be used either as an alternative to the metabolic conversion of plastic to CO_2_, or as the initial oxidative step in combination with standard microbial degradation pathways.

Further on, this study suggests that insect saliva might result as a depository of degrading enzymes (plastic, cellulose or lignin to mention some) which could revolutionize the bioremediation field. Although further studies will be necessary to obtain a deeper understanding of the step-by-step evolution of plastic in contact with ww saliva enzymes, this discovery introduces a new paradigm for dealing with plastic degradation. In a circular economy frame, this study opens up a new potential field both in plastic upcycling, and in manufacturing the plastic of the future, with ad-hoc formulations prone to facilitate degradation by selected enzymes.

## Acknowledgments

We thank the Proteomics and Genomics, Electron Microscopy and Gas Chromatography facilities of the CIB for the excellent technical support. We thank Paloma Delgado and Pedro Velasco from InsectPark-Microfauna S.L., El Escorial, Madrid, Spain (https://insectpark.es/) for providing *Samia cynthia* larvae. We thank the Centres Científics i Tecnolo`gics Unitat Cromatografia de Gases-Espectrometria de Masses Aplicada (CCIT), University of Barcelona, Spain for the support provided in the GC-MS experiments. We thank the Roechling foundation for supporting and sponsoring this work.

## Funding

Roechling Stiftung to FB

Consejo Superior de Investigaciones Científicas (CSIC) to FB

NATO Science for Peace and Security Programme (Grant SPS G5536) to TT

Junta de Castilla y León, Consejería de Educación y Cultura y Fondo Social Europeo (Grant BU263P18) to TT

Ministerio de Ciencia e Innovación (Grant PID2019-111215RB-100) to TT The Generalitat de Catalunya (2017 SGR 1192) to MS

Ministerio de Ciencia e Innovación (Grant BFU2017-89143-P) to EA-P

## Author contributions

Conceptualization: FB, CFA

Methodology: All the Authors

Investigation: All the authors

Visualization: FB, CFA, PCV, LG, MS, EA-P

Funding acquisition: FB, TT, MS, EA-P

Project administration: FB

Writing original draft: FB

Writing, review & editing: All the authors

## Methods

### Resources availability

Further information and requests for resources and reagents should be directed to and will be fulfilled by the lead contact, Federica Bertocchini.

### Materials availability

This study did not generate new unique reagents.

### Data availability

The RNA-seq data that was used for the *Galleria mellonella* genome annotation has been deposited in NCBI under the BioProject accession number PRJNA822887. In addition, the newly generated annotation in GFF3/GTF format was deposited to Data Mendeley (doi:10.17632/t7b5s58vxt.2).

### Experimental model and subject details

*Galleria mellonella* larvae colony is maintained in an incubator at 28 °C in the dark, and fed with beeswax from beehives.

### Wax worm saliva collection

Larvae of 150-300 mg were used for saliva collection. Briefly, a glass capillary connected to a mouth pipet was placed at the buccal opening and the liquid was collected. Saliva for PE application was immediately used. Occasionally, frozen saliva-only can be utilized. For electron microscopy, saliva was diluted 1:1 in the following buffer: 10mM Tris-Cl pH 8, 50 mM NaCl. For proteomic analyses, saliva was diluted 1:1 in 10 mM Tris-Cl, 50 mM NaCl, 2 mM DTT, 20% glycerol. For control, *Samia cynthia* larvae (kindly provided by InsectPark-Microfauna S.L. (Escorial, Madrid) at the last stage were used to collect saliva as previously described.

### RAMAN and FTIR analyses

PE film was treated with 30 *μ*l of GmSal for 90 minutes, three applications for RAMAN. PE film was treated with 5 *μ*l of GmSal for 90 minutes, nine applications for FTIR. Recombinant proteins were applied as follows: 5 *μ*l of protein (concentration between 1 and 5 *μ*g/ml) were applied eight times on PE film 90 minutes each time. For the control with inactivated proteins, recombinant proteins were denatured at 100 degrees for 10 minutes. SEC and IEX peak aliquots were applied six times, 30 minutes each, and left overnight. Treated and control films were washed with water and ethanol. RAMAN analyses were performed on (treated and control) PE films using Alpha300R –Alpha300A AFM Witec equipment with 5mW power, 50x (NA0.8) objective, integration time 1, accumulation 30, wavelength 532nm. FTIR analyses were performed with a Jasco LE-4200 equipment, with the following features: interval 4000-400 cm^−1^, Resolution 4 cm^−1^, scan 264.

### High Temperature-Gel Permeation Chromatography (HT-GPC) analysis

HT-GPS was performed by Polymer Chart, Valencia, Spain. Briefly, the followings are the experimental conditions: Equipment: GPC-IR5 I Polymer Char; solvent: TCB stabilized with 300ppm of BHT; dissolution temperature, detectors temperature, columns Temperature: 160 degrees; volume: 8ml; weight: 8mg; dissolution Time: 60 minutes; injected volume: 200 *μ*l; injection time: 55 minutes; flow: 1ml/min; columns: 3 × PL gel Olexis Mix-Bed columns (13 microns), 300 × 7.5mm + guard column For the carbonyl index analysis, GPC IR6 was used, with dissolvent o-DCB and temperature at 150 degrees.

PE film and PE 4000 were treated with 100 *μ*l of GmSal for 90 minutes. The treatment was repeated 15 times (film and PE 4000) and 30 times (PE 4000).

### Gas Chromatography Mass Spectroscopy (GC-MS) and Tandem analysis

An amount of 20 mg of PE 4,000 or 1.5 mg PE 2,000 were placed in a 1.5 ml Eppendorf tube. PE was exposed to 40 *μ*l of *G. mellonella* saliva 9 times for 90 minutes each at room temperature and avoiding light. For prolonged treatment (day 1, 2, 3, and 6), three applications of 100 *μ*l of saliva for 90 minutes each at room temperature were carried out each day. Controls of each PE were performed using Milli-Q water in substitution of the saliva of *G. mellonella* larvae, as well as saliva of *G. mellonella* larvae only. Also, PE was exposed to 10 *μ*l (1.2 mg/mL) of Demetra 24 times for 90 minutes. Prolonged treatment was performed as well for Demetra (days 1 and 2), five applications per day of 10 *μ*l (1.2 mg/mL) for 90 minutes each.

As control, the same experiment was repeated using the protein buffer. Afterward, samples were centrifuged with an Eppendorf centrifuge 5810 R at 19,083 G-force for 30 seconds and the subnatant was transferred to a new 1.5 ml Eppendorf tube. Samples and controls were extracted using a QuEChERS (quick, easy, cheap, effective, and safe) method [1] based on [2] with some modifications. Briefly, 50 *μ*l of diphenyl phthalate (Internal Standard; IS) at a concentration of 1 mg/ml was added at each sample and extracted with 300 *μ*l of dichloromethane (DCM) and 5 % (v/m) of NaCl. The tube was vortexed for 30 seconds and sonicated in a bath (50/60 Hz) for 15 min at room temperature, followed by centrifugation with an Eppendorf centrifuge 5810 R at 20 ^*°*^C and 19,083 G-force for 10 min. Finally, DCM located as the subnatant was collected and placed in an insert before analysis. Silylation reaction with N, O-Bis(trimethylsilyl)trifluoroacetamide (BSTFA) was performed to determine the low-volatility polar compounds which show low detection sensibility. A fraction of 50 *μ*l of each sample with 50 *μ*l of BSTFA was incubated for 20 min at 60 ^*°*^C before the analysis.

Dichloromethane (DCM; CAS-No: 75-09-2) for gas chromatography-mass spectrometry (GC-MS) was SupraSolv grade purity and obtained from Sigma-Aldrich (Darmstadt, Germany). Sodium chloride (NaCl; ≥ 99.5%; CAS-No: 7647-14-5) and ultrapure water from a Milli-Q system were supplied from Merck (Darmstadt, Germany).

Crystalline granular powder polyethylene (PE 4000; CAS-No: 9002-88-4, specification sheet available at https://www.sigmaaldrich.com/ES/es/product/aldrich/427772) was supplied by Sigma-Aldrich (Saint Louis, USA).

Chromatographic analyses were performed with a gas chromatography-mass spectrometry system (GC-MS) 7980A-5975C from Agilent Technologies. Separation of the metabolites was performed on a DB-5th Column coated with polyimide (30 m length, 0.25 mm inner diameter, and 0.1 *μ*m film thickness; Agilent Technologies, USA) for proper separation of substances, and Helium (He) was utilized as a carrier gas. The analysis was performed using a split injector at 350 ^*°*^C and an injection volume of 1 *μ*l. The ion source temperature was 230 °C, the ^*°*^C mass spectral analysis was performed in scan mode, the quadrupole temperature of 150 ^*°*^C, and a fragmentation voltage of 70 eV. The oven program started at 60 ^*°*^C for 3 min, then 20 ^*°*^C/min to 350 ^*°*^C for 1 min. The total run time was 18.5 min and 19.5 min for derivatized samples. The resulting chromatograms were processed using the software MSD ChemStation E.01.00.237 from Agilent Technologies, Inc while for the identification NIST11 library was used.

The evaluation of the prolonged treatment was based on the relative abundance of each untargeted compound, which consists of the quotient of the area under the peak of each compound divided by the area under the peak of the IS.

Gas Chromatography/Tandem Mass Spectrometry was used for confirmation of the non-target compounds by a BRUKER 456-GC SCION TQ. This experiment was performed by the Elemental and Molecular Analysis facility, University of Extremadura, Spain. Briefly, the injector port was set at 230 ^*°*^C in Split mode. Separation was achieved using a column HP 5MS, 30 m, 0.25 mm, and 0.25 *μ*m. Helium (He) was utilized as a carrier gas. The column oven was programmed in the following conditions: 60 ^*°*^C for 3 min, increase of 20 ^*°*^C/min to 325 °C for 1 min. The collision energy was 15 eV.

### Electron Microscopy analysis

Larvae saliva samples were diluted 1:50 in the proper buffer (see section “Wax worm saliva collection”). Samples were analyzed by electron microscopy (EM) after being adsorbed to glow-discharged carbon coated grids and stained with 2% uranyl acetate. Grids were observed using a JEOL JEM-1230 EM operated at 100 kV and a nominal magnification of 40 000. EM images were taken under low dose conditions with a CMOS Tvips TemCam-F416 camera, at 2.84 Å per pixel.

### Protein Chromatography analyses

For the size exclusion chromatography, ww saliva in the proper buffer (see “Wax worm saliva collection”) was thawed, pooled and centrifuged. The supernatant was filtered (0.45 *μ*m cutoff, Ultrafree Millipore) and loaded to a size exclusion chromatography column Superdex 200 5-150 (Cytiva) equilibrated with 10 mM Tris-Cl, 50 mM NaCl, 2 mM DTT. For the ion exchange chromatography, upon thawing, the sample was diluted to 100 *μ*l with 10 mM Tris-Cl at pH 8, centrifuged, filtered and the supernatant loaded to a monoQ 5/50 GL ion exchange column (Cytiva). After a wash step, a 40ml gradient with buffer A (10 mM Tris-Cl pH8), and buffer B (same as A supplemented with 500 mM NaCl) was applied.

### Proteomic analysis

Liquid Chromatography Mass Spectrometry (LC-MS) analysis. All peptide separations were carried out on an Easy-nLC 1000 nano system (Thermo Fisher Scientific). For each analysis, the sample was loaded into a precolumn Acclaim PepMap 100 (Thermo Fisher Scientific) and eluted in a RSLC PepMap C18, 15 cm long, 50 *μ*m inner diameter and 2 *μ*m particle size (Thermo Fisher Scientific). The mobile phase flow rate was 300 nl/min using 0.1% formic acid in water (solvent A) and 0.1% formic acid and 100% acetonitrile (solvent B). The gradient profile was set as follows: 5%–35% solvent B for 45 min, 35%-100% solvent B for 5 min, 100% solvent B for 10 min. Four *μ*l of each sample were injected.

MS analysis was performed using a Q Exactive mass spectrometer (Thermo Fisher Scientific). For ionization, 1900 V of liquid junction voltage and 270 ^*°*^C capillary temperature was used. The full scan method employed a m/z 400–1500 mass selection, an Orbitrap resolution of 70,000 (at m/z 200), a target automatic gain control (AGC) value of 3e6, and maximum injection times of 100 ms. After the survey scan, the 15 most intense precursor ions were selected for MS/MS fragmentation. Fragmentation was performed with a normalized collision energy of 27 eV and MS/MS scans were acquired with a starting mass of m/z 100, AGC target was 2e5, resolution of 17,500 (at m/z 200), intensity threshold of 8e4, isolation window of 2 m/z units and maximum IT was 100 ms. Charge state screening was enabled to reject unassigned, singly charged, and equal or more than seven protonated ions. A dynamic exclusion time of 20s was used to discriminate against previously selected ions.

MS data analysis. Mass spectra *.raw files were searched against an in –house specific database against Galleria proteins (12715 proteins entries), using the Sequest search engine through Proteome Discoverer (version 1.4.1.14) (Thermo Scientific). Search parameters included a maximum of two missed cleavages allowed, carbamidomethyl of cysteines as a fixed modification and oxidation of methionine as variable modifications. Precursor and fragment mass tolerance were set to 10 ppm and 0.02 Da, respectively. Identified peptides were validated using Percolator algorithm with a q-value threshold of 0.01. The protein identification by nLC-MS/MS was carried out in the Proteomics and Genomics Facility (CIB-CSIC), a member of ProteoRed-ISCIII network [3].

### Recombinant protein production and utilization

Arylphorin, Arylphorin subunit alpha-like (Demetra) and hexamerin (Ceres) were produced by Genscript, utilizing the baculovirus expression system in insect cells, according to the manufacturer. Briefly, sf9 cells were infected with P2 baculovirus, flasks were incubated at 27 ^*°*^C for 48-72 h and media harvested. Then cells were removed, and transfection medium was applied for purification. The produced proteins were resuspended in 150 mM NaCl, 20 mM Hepes, 5% glycerol and used for the degradation assay. The same buffer alone was used as negative control.

### *Galleria mellonella* genome annotation

In order to obtain the most useful information from mass spectrometry analysis of proteins extracted from ww saliva, it is pivotal to use a representative database of protein sequences. In addition to the NCBI official annotation of *Galleria mellonella* protein sequences, a new annotation was produced exploiting also the information of *G. mellonella* salivary glands RNA-seq data. *G. mellonella* genome annotation was performed using the genome sequence available at NCBI. A specific pipeline was developed to combine the information from RNA-seq with *ab initio* predictors in order to obtain the most accurate annotation. Briefly, the RNA-seq data was mapped on the reference genome using STAR (version 2.5 0c) [4] in local mode and used to perform a reference guided transcriptome assembly with Trinity (v2.11.0) [5]. The obtained transcripts and the mapping files were used as input for the Braker2 pipeline [6] to combine AUGUSTUS *ab initio* annotation [7] with the transcriptome assembly to obtain the annotation in GFF format, together with transcript and protein sequences. Proteins were used as input for the PANNZER2 pipeline to obtain descriptions [8], Gene Ontology and KEGG annotations. About 32000 genes could be annotated in the *Galleria genome*. The corresponding proteins were analyzed to assess their completeness performing a BLASTP alignment against the UniRef90 database and calculating the percentage of alignment. A similarity search against the UniRef90 (Nov2018) database showed that about 50% of the predicted proteins covered 100% of the corresponding hits (i.e. full length) and that about 80% of the predicted proteins covered at least 50% of the corresponding hits. As a further control, the proteins and the genome were evaluated with the BUSCOv2 pipeline [9]. The BUSCO database contains sets of single-copy highly conserved genes across different taxa (i.e. Eukaryota or Insects). By performing an analysis with the BUSCO database it is possible to assess the completeness of a genome/proteome, the presence of duplications and/or fragmentations. This analysis was performed using the predicted proteome and also the unannotated genome for a comparison. By comparing the results of the unannotated with the annotated genome, we can see a small fraction of missing genes which are probably absent from the genome assembly and that cannot be recovered from the current genome sequence. This explains some missing genes present in the later NCBI annotation.

The RNA-seq data that was used for the G. mellonella genome annotation has been deposited in NCBI under the BioProject accession number PRJNA822887. In addition, the newly generated annotation in GFF3/GTF format was deposited in Data Mendeley (doi:10.17632/t7b5s58vxt.2).

## Supplementary Information

**Figure S1.**
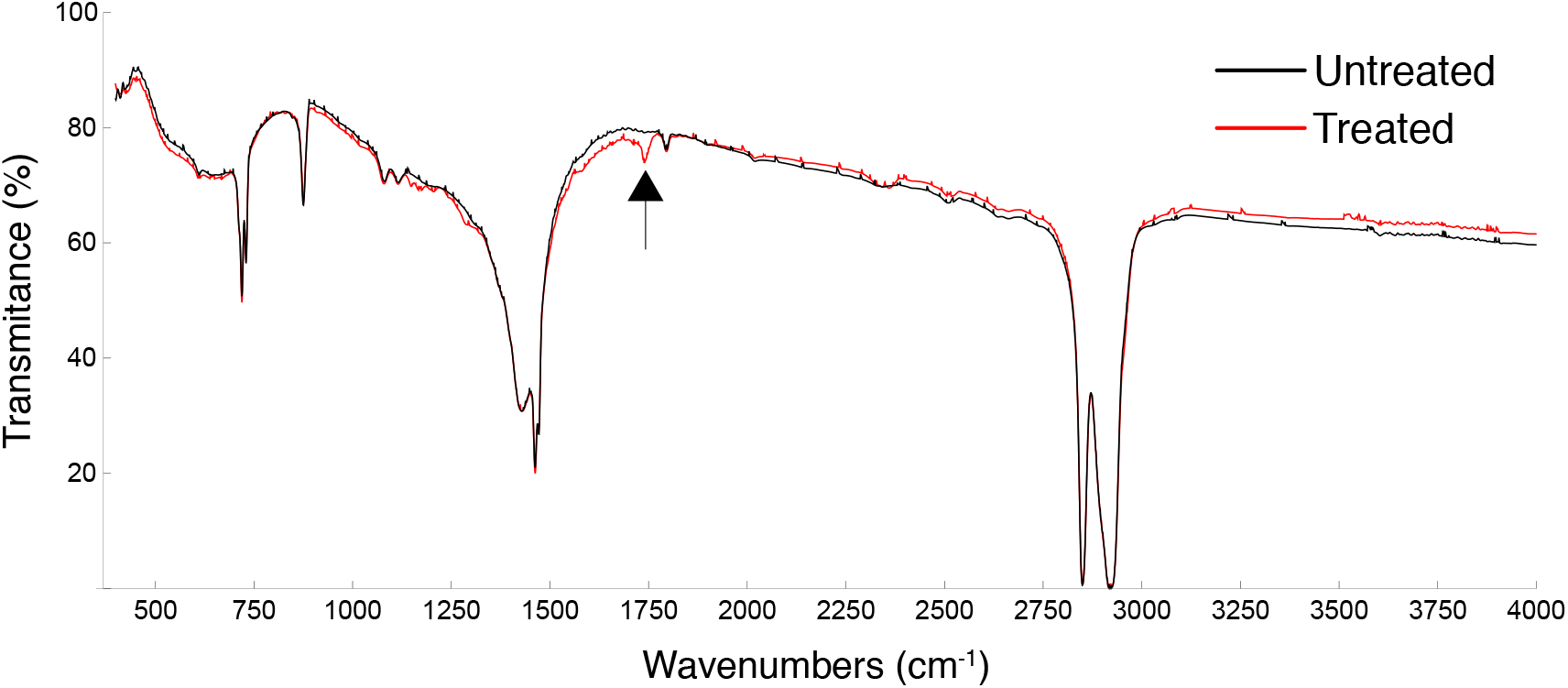
FTIR of PE film treated with GmSal. The arrow indicates the peak at around 1750 cm^−1^ (i.e. oxidation-carbonyl group).

**Figure S2.**
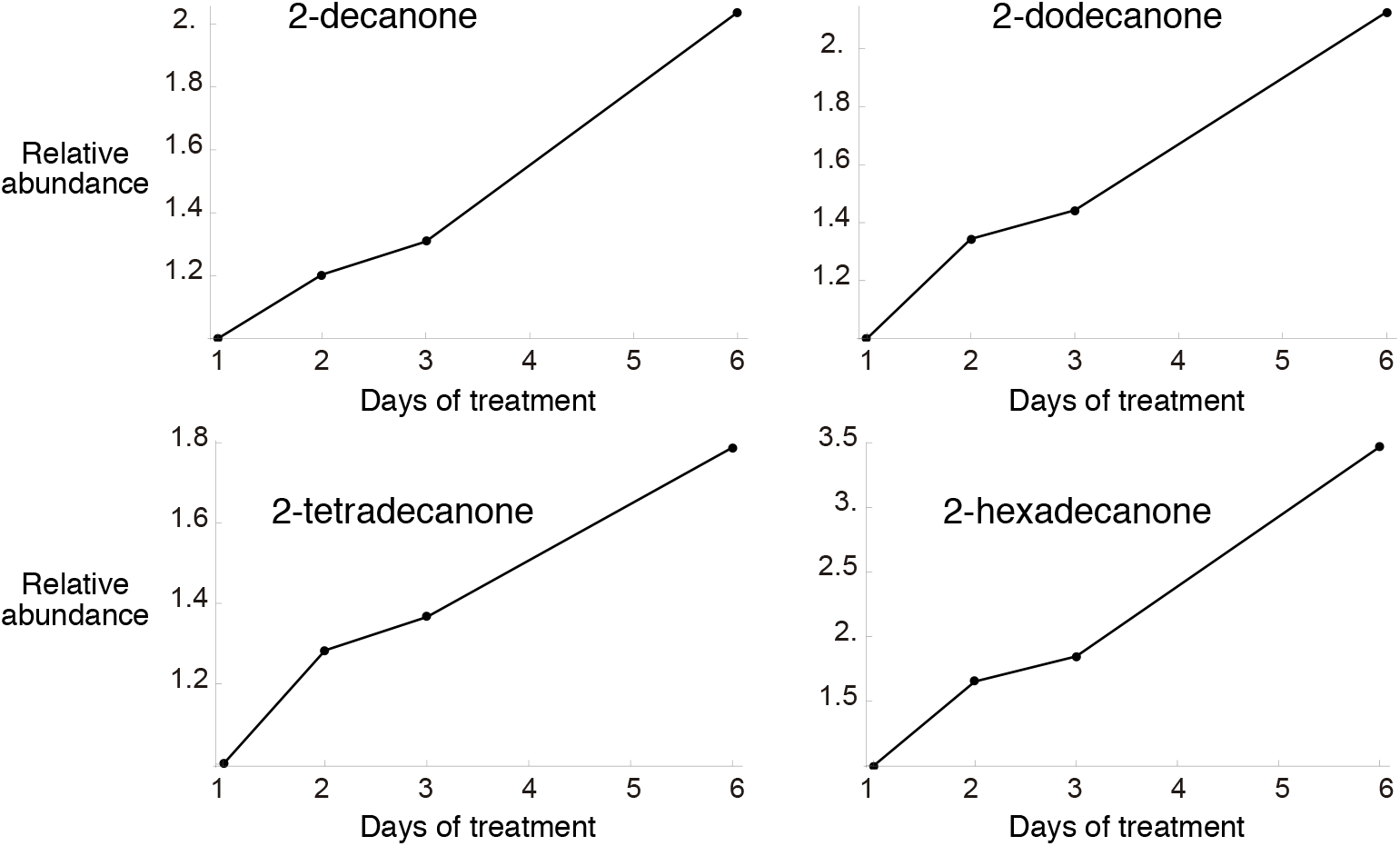
Relative abundance of the ketones 2-decanone, 2-dodecanone, 2-tetradecanone, 2-hexadecanone measured at day 1, 2, 3 and 6 of PE treatment with GmSal.

**Figure S3.**
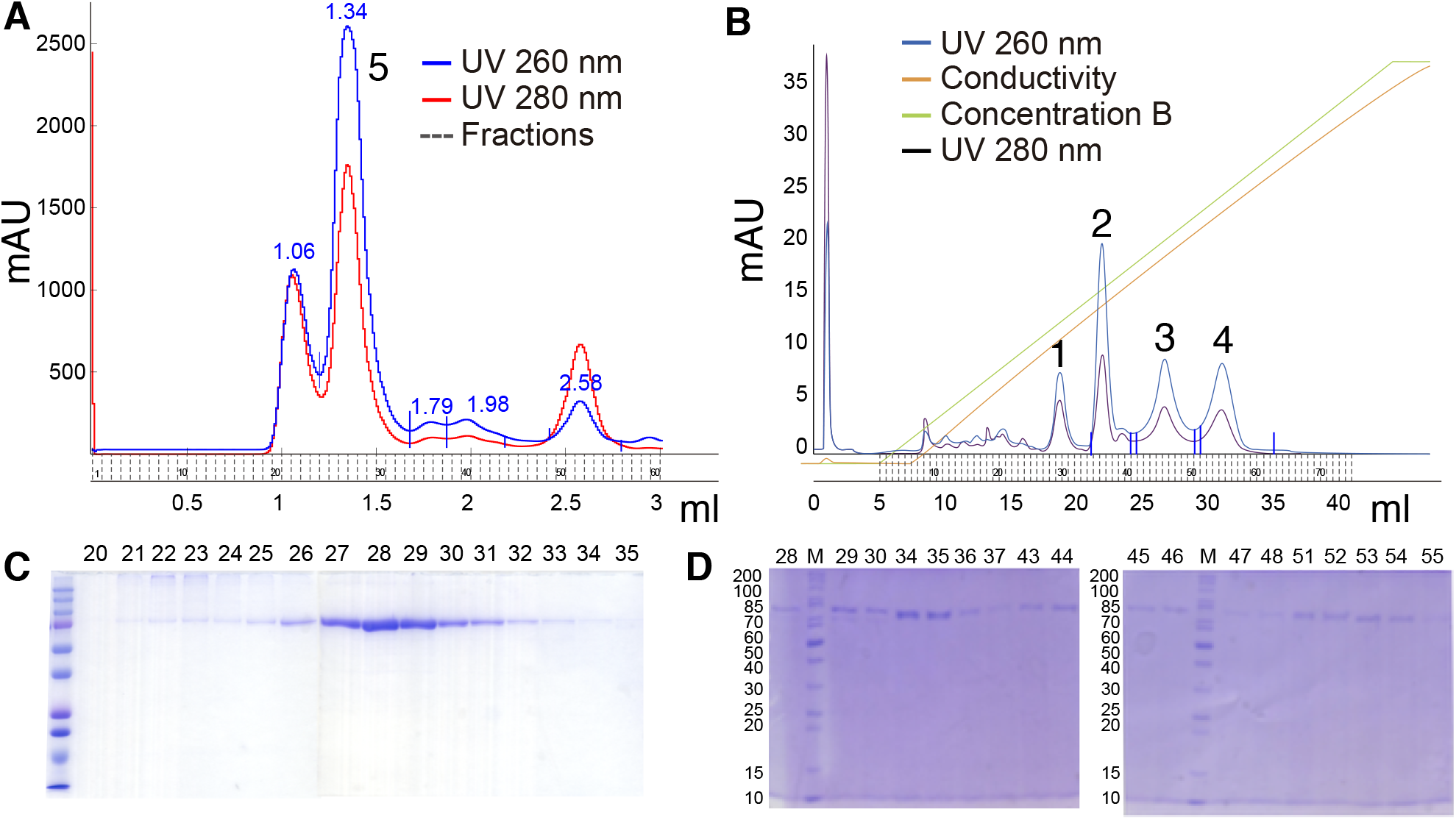
Chromatographic analyses of saliva samples. A, B. Size exclusion chromatography (A), and ion exchange chromatography (B). C. SDS-gel of the fractions in A. D. SDS-gel of the fractions in B.

**Figure S4.**
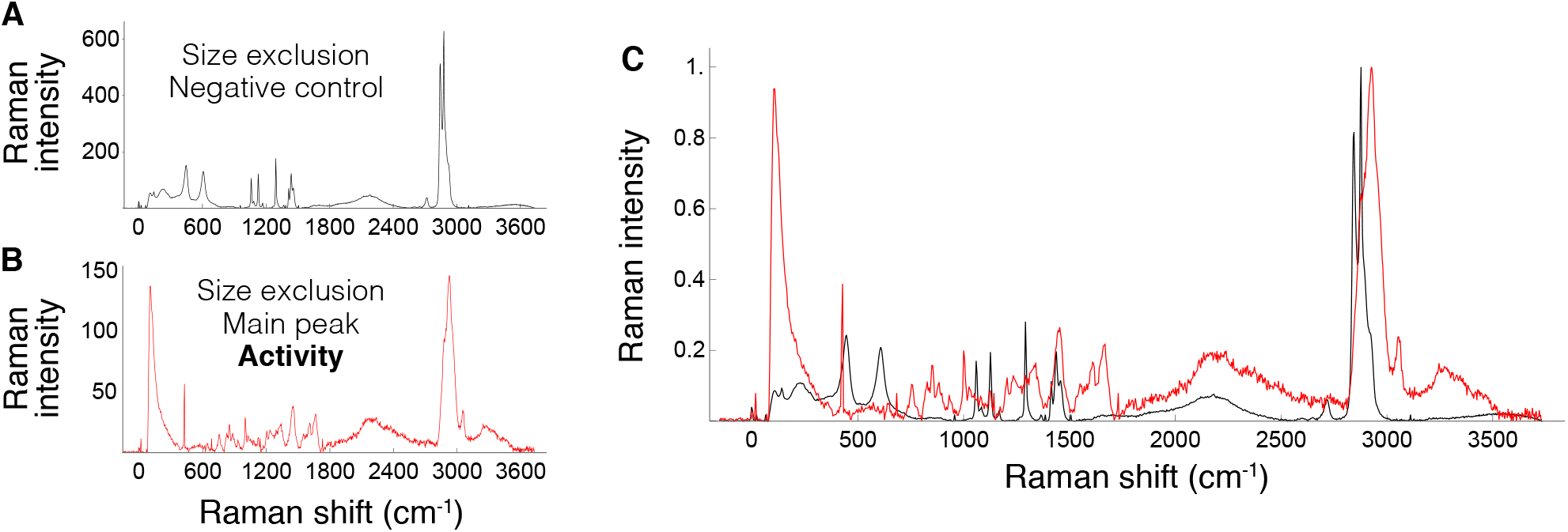
Functional analysis of degradation activity in the size exclusion main fraction (peak 5 in Fig. S3A). A, B AFM-RAMAN spectroscopy of control (A) and peak 5-treated PE film (B). A. Control PE film (see Fig. 1 for details). B. Punctual analyses of treated PE film, indicating PE deterioration. The spectrum shows an intense peak below 600 cm^−1^, indicating an increase in additive detection; peaks at 711 and 1747 cm^−1^ correspond to the C=O group; oxidation also indicated between 3000-3500 cm^−1^, and PE deterioration revealed by the broad peak between 1500 and 2400 cm^−1^. C. Overlapping of A and B.

**Figure S5.**
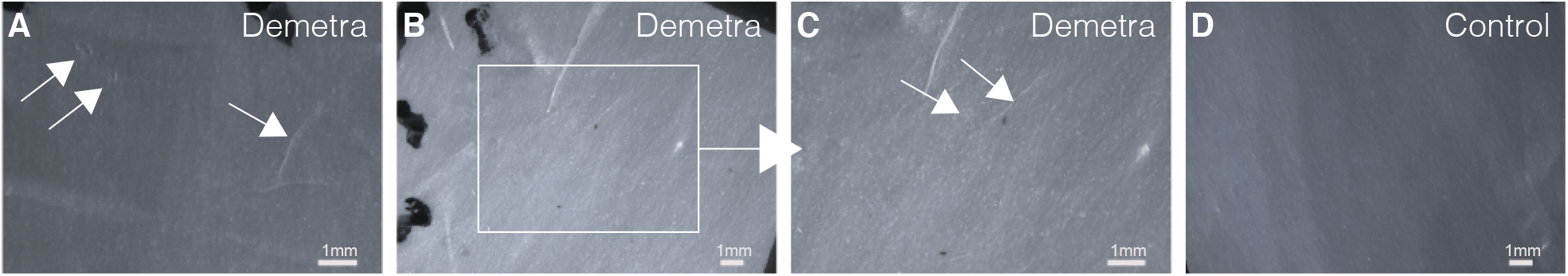
Demetra effect on PE film. A-D. Demetra causes diverse extent of damage after application on PE film, with some visible external signs, (A and magnification in B) to milder, barely visible effects (C, arrows). D. Control.

**Figure S6.**
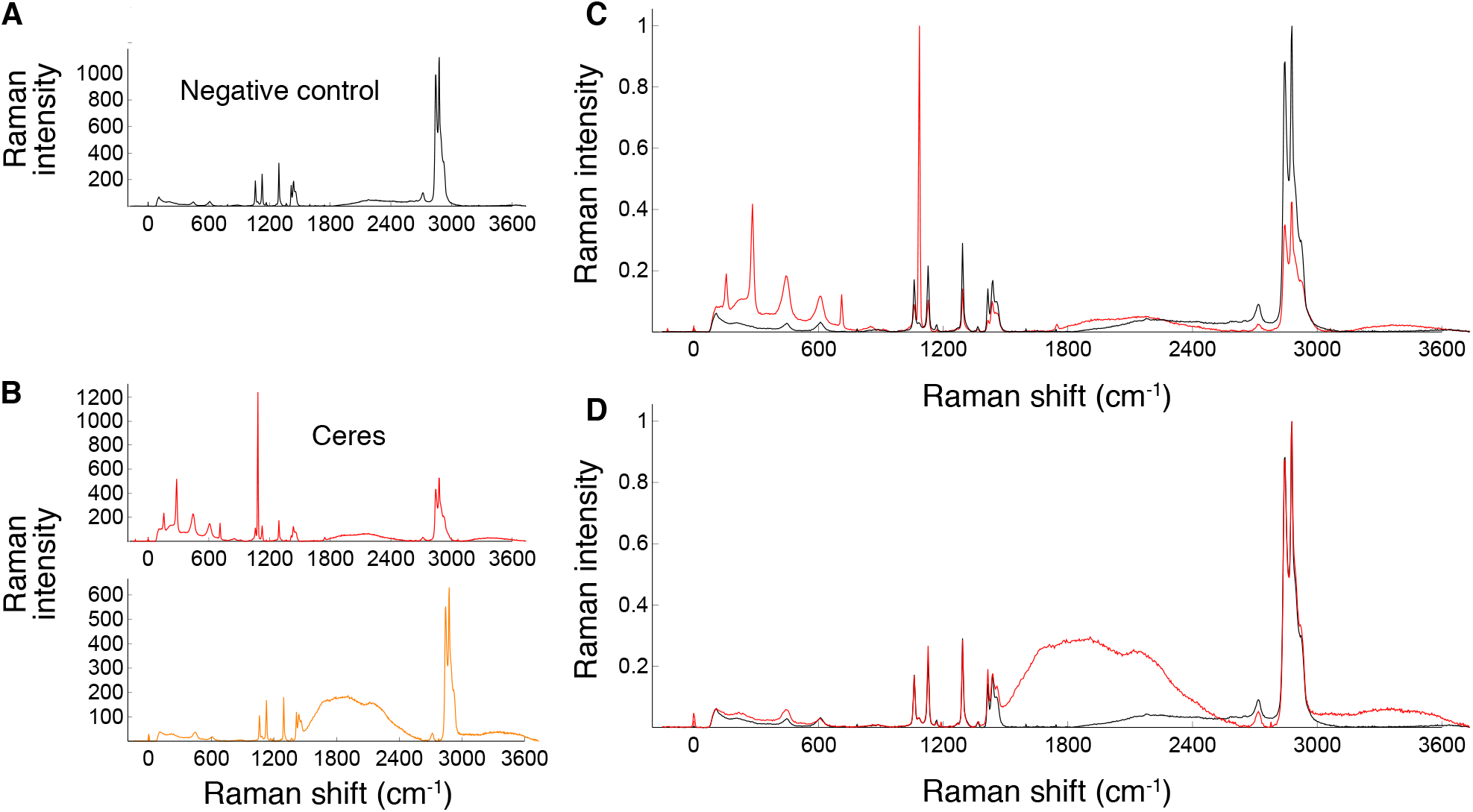
Ceres effect on PE film. A, B AFM-RAMAN spectroscopy of control (A) and Ceres treated PE film (B, C). A. Control PE film, showing the typical PE peaks at 1061, 1128, 1294, 1440, 2846 and 2880 cm^−1^. B, C. Punctual analyses of treated PE film, indicating PE deterioration. The spectrum shows an intense peak at 1085 cm^−1^ assigned to amorphous PE (B). The peaks at 711 and 1747 cm^−1^ correspond to the C=O group. D, E. Overlapping of the negative control with the experimental sample in B (D), and C (E). The broad band between 1500 and 2400 cm^−1^ (B, C) indicates different collective stretching vibrations due to the presence of other organic compounds, that is PE deterioration.

**Figure S7.**
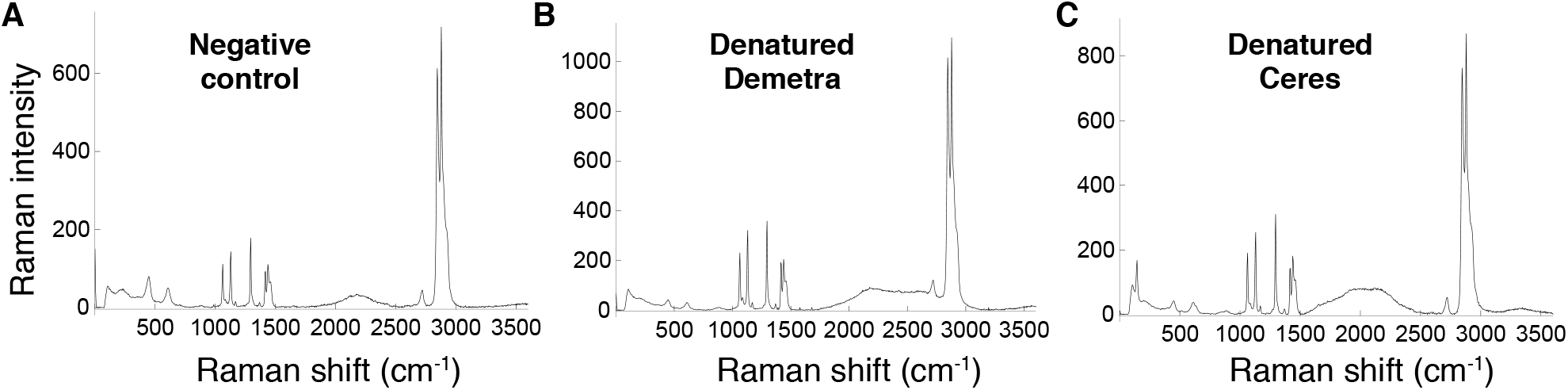
Denatured Demetra and Ceres on PE film. A-C. RAMAN spectroscopy of control PE (A) and PE treated with denatured Demetra (B) and Ceres (C).

**Table S1.** Molecular weight (uma), fragment ions (m/z), and intensity of the untargeted detected compounds from the NIST Mass Spectrometry library compared to those detected in our study.

| Molecule | Molecular weight (uma) | Mass spectrum (NIST Mass Spectrometry library) |  |  |  |  | Mass spectrum of the sample |  |  |  |  |
| --- | --- | --- | --- | --- | --- | --- | --- | --- | --- | --- | --- |
|  |  | m/z (intensity) |  |  |  |  | m/z (intensity) |  |  |  |  |
| 2-decanone (10 C) | 156 | 58 (99.9) | 43 (84.5) | 71 (37.7) | 59 (27.9) | 41 (27.3) | 58 (99.9) | 43 (92.0) | 41 (51.2) | 44 (44.7) | 55 (44.3) |
| 2-dodecanone (12 C) | 184 | 58 (99.9) | 43 (93.4) | 71 (38.7) | 59 (34.4) | 41 (32.3) | 58 (99.9) | 43 (46.1) | 59 (34.3) | 41 (22.7) | 71 (22.1) |
| 2-tetradecanone (14 C) | 212 | 58 (99.9) | 43 (96.3) | 59 (43.1) | 71 (34.6) | 41 (31.6) | 58 (99.9) | 43 (61.6) | 59 (38.3) | 71 (36.5) | 41 (26.6) |
| 2-hexadecanone (16 C) | 240 | 58 (99.9) | 59 (61.9) | 43 (59.4) | 71 (39.3) | 41 (25.8) | 58 (99.9) | 43 (86.8) | 59 (59.7) | 71 (41.2) | 41 (30.4) |
| 2-octadecanone (18 C) | 268 | 58 (99.9) | 59 (75.2) | 43 (59.7) | 71 (30.7) | 41 (23.7) | 43 (99.9) | 58 (87.7) | 71 (65.8) | 59 (54.4) | 41 (51.3) |
| 2-eicosanone (20 C) | 296 | - | - | - | - | - | 58 (99.9) | 43 (82.7) | 71 (73.4) | 41 (69.3) | 55 (64.6) |
| 2-docosanone (22 C) | 324 | - | - | - | - | - | 58 (99.9) | 59 (93.8) | 43 (75.9) | 57 (67.8) | 44 (6.22) |
| 2,3-Butanediol, 2TMS derivative | 234 | 117 (99.9) | 73 (83.3) | 147 (34.2) | 75 (15.1) | 118 (11.0) | 117 (99.9) | 73 (59.2) | 147 (37.5) | 75 (9.4) | 118 (9.0) |
| Benzenepropanoic acid, TMS derivative | 222 | 104 (99.9) | 75 (85.0) | 73 (47.0) | 207 (41.5) | 91 (27.0) | 104 (99.9) | 75 (80.2) | 73 (47.8) | 207 (36.1) | 91 (30.0) |
| Sebacic acid, 2TMS derivative | 346 | 331 (99.9) | 73 (87.4) | 75 (78.0) | 215 (49.7) | 129 (38.1) | 331 (99.9) | 215 (78.2) | 73 (67.4) | 75 (40.3) | 129 (16.5) |

## References

1. https://plasticseurope.org/

2. Ellis, L.D., Rorrer, N. A., Sullivan, K. P., Otto, M., McGeehan, J. E., Roman-Leshkov, Y., Wierckx, N. and Beckham, G. T. (2021). Chemical and biological catalysis for plastics recycling and upcycling. Nature Catalysis 4, 539–556. 10.1038/s41929-021-00648-4

3. Zhang, F., Wang, F., Wei, X., Yang, Y., Xu, S., Deng, D. and Wang, Y.-Z. (2022). From trash to treasure: Chemical recycling and upcycling of commodity plastic waste to fuels, high-valued chemicals and advanced materials. Journal of Energy Chemistry 69, 369–388. 10.1016/j.jechem.2021.12.052

4. https://goldbook.iupac.org/terms/view/B00656

5. Vert, M., Doi, Y., Hellwich, K.-H., Hess, M., Hodge, P., Kubisa, P., Rinaudo, M. and Schué, F. (2012). Terminology for biorelated polymers and applications (IUPAC Recommendations 2012). Pure and Applied Chemistry 84, 377–410. 10.1351/PAC-REC-10-12-04

6. Albertsson, A.-C.a.K. S. (1990). The influence of biotic and abiotic environments on the degradation of polyethylene. Progress in Polymer Science 15, 177–192. 10.1016/0079-6700(90)90027-X

7. Gryn’ova, G., Hodgson, J.L., and Coote, M.L. (2011). Revising the mechanism of polymer autooxidation. Org Biomol Chem 9, 480–490. 10.1039/c0ob00596g

8. Albertsson, A.-C., Andersson, S. O. and Karlsson, S. (1987). The mechanism of biodegradation of polyethylene. Polymer Degradation and Stability 18, 73–87. 10.1016/0141-3910(87)90084-X

9. Roy, P.K., Hakkarainen, M., Varma, I.K., and Albertsson, A.C. (2011). Degradable polyethylene: fantasy or reality. Environ Sci Technol 45, 4217–4227. 10.1021/es104042f

10. Hakkrainen, M.a.A. A.-C. (2004). Environmental degradation of polyethylene. Adv Polym Sci 169, 177–199. 10.1007/b13523

11. Koutny, M., Lemaire, J., and Delort, A.M. (2006). Biodegradation of polyethylene films with prooxidant additives. Chemosphere 64, 1243–1252 10.1016/j.chemosphere.2005.12.060

12. Wei, R.a.Z. W. (2017). Microbial enzymes for the recycling of recalcitrant petroleum-based plastics: how far are we? Microbial Biotechnology 10, 1308–1322. 10.1111/1751-7915.12710

13. Restrepo-Florez, J.-M., Bassi, A. and Thompson, M. R. (2014). Microbial degradation and deterioration of polyethylene. International Biodeterioration & Biodegradation 88, 83–90. 10.1016/j.ibiod.2013.12.014

14. Amobonye, A., Bhagwat, P., Singh, S., and Pillai, S. (2021). Plastic biodegradation: Frontline microbes and their enzymes. Sci Total Environ 759, 143536. 10.1016/j.scitotenv.2020.143536

15. Matjašič, T., Simčič, T., Medvešček, N., Bajt, O., Dreo, T. and Mori, N. (2021). Critical evaluation of biodegradation studies on synthetic plastics through a systematic literature review. Science of the Total Environment 752. 10.1016/j.scitotenv.2020.141959

16. Walsh, A.N., Reddy, C. M., Niles, S. F., McKenna, A. M., Hansel, C. M. and Ward, C. P. (2021). Plastic formulation is an emerging control of its photochemical fate in the ocean. Environ. Sci. Technol 55, 12383–12392. 10.1021/acs.est.1c02272

17. Peixoto, J., Silva, L.P., and Kruger, R.H. (2017). Brazilian Cerrado soil reveals an untapped microbial potential for unpretreated polyethylene biodegradation. J Hazard Mater 324, 634–644. 10.1016/j.jhazmat.2016.11.037

18. Yoon, M.G., Jeon, H. J. and Kim, M. N. (2012). Biodegradation of polyethylene by a soil bacterium and AlkB cloned recombinant cell. Journal of Bioremediation & Biodegradation 3. 10.4172/2155-6199.1000145

19. Kyaw, B.M., Champakalakshmi, R., Sakharkar, M.K., Lim, C.S., and Sakharkar, K.R. (2012). Biodegra-dation of low density polythene (LDPE) by Pseudomonas Species. Indian J Microbiol 52, 411–419. 10.1007/s12088-012-0250-6

20. Dey, A.S., Bose, H., Mohapatra, B., and Sar, P. (2020). Biodegradation of unpretreated low-density polyethylene (LDPE) by Stenotrophomonas sp. and Achromobacter sp., isolated from waste dumpsite and drilling fluid. Front Microbiol 11, 603210. 10.3389/fmicb.2020.603210

21. Delacuvellerie, A., Cyriaque, V., Gobert, S., Benali, S., and Wattiez, R. (2019). The plastisphere in marine ecosystem hosts potential specific microbial degraders including Alcanivorax borkumensis as a key player for the low-density polyethylene degradation. J Hazard Mater 380, 120899. 10.1016/j.jhazmat.2019.120899

22. Mohanan, N., Montazer, Z., Sharma, P.K., and Levin, D.B. (2020). Microbial and enzymatic degradation of synthetic plastics. Front Microbiol 11, 580709. 10.3389/fmicb.2020.580709

23. Montazer, Z., Habibi Najafi, M.B., and Levin, D.B. (2020). Challenges with verifying microbial degradation of polyethylene. Polymers (Basel) 12. 10.3390/polym12010123

24. Santo, M., Weitsman, R. and Sivan, A. (2013). The role of the copper-binding enzyme – laccase – in the biodegradation of polyethylene by the actinomycete Rhodococcus ruber. International Biodeterioration & Biodegradation 84, 204–2010. 10.1016/j.ibiod.2012.03.001

25. Fujisawa, M., Hirai, H. and Nishida, T. (2001). Degradation of polyethylene and nylon-66 by the laccase-mediator system. Journal of Polymers and the Environment 9, 103–108. 10.1023/A:1020472426516

26. Yang, J., Yang, Y., Wu, W.M., Zhao, J., and Jiang, L. (2014). Evidence of polyethylene biodegradation by bacterial strains from the guts of plastic-eating waxworms. Environ Sci Technol 48, 13776–13784. 10.1021/es504038a

27. Yang, Y., Yang, J., Wu, W.M., Zhao, J., Song, Y., Gao, L., Yang, R., and Jiang, L. (2015). Biodegradation and mineralization of polystyrene by plastic-eating mealworms: Part 2. Role of gut microorganisms. Environ Sci Technol 49, 12087–12093. 10.1021/acs.est.5b02663

28. Yang, Y., Yang, J., Wu, W.M., Zhao, J., Song, Y., Gao, L., Yang, R., and Jiang, L. (2015). Biodegradation and mineralization of polystyrene by plastic-eating mealworms: Part 1. chemical and physical characterization and isotopic tests. Environ Sci Technol 49, 12080–12086. 10.1021/acs.est.5b02661

29. Bombelli, P., Howe, C.J., and Bertocchini, F. (2017). Polyethylene bio-degradation by caterpillars of the wax moth Galleria mellonella. Curr Biol 27, R292–R293. 10.1016/j.cub.2017.02.060

30. Brandon, A.M., Gao, S.H., Tian, R., Ning, D., Yang, S.S., Zhou, J., Wu, W.M., and Criddle, C.S. (2018). Biodegradation of polyethylene and plastic mixtures in mealworms (larvae of Tenebrio molitor) and effects on the gut microbiome. Environ Sci Technol 52, 6526–6533. 10.1021/acs.est.8b02301

31. Yang, Y., Wang, J., and Xia, M. (2020). Biodegradation and mineralization of polystyrene by plastic-eating superworms Zophobas atratus. Sci Total Environ 708, 135233. 10.1016/j.scitotenv.2019.135233

32. Kong, H.G., Kim, H.H., Chung, J.H., Jun, J., Lee, S., Kim, H.M., Jeon, S., Park, S.G., Bhak, J., and Ryu, C.M. (2019). The Galleria mellonella hologenome supports microbiota-independent metabolism of long-chain hydrocarbon beeswax. Cell Rep 26, 2451–2464 e2455. 10.1016/j.celrep.2019.02.018

33. Peydaei, A., Bagheri, H., Gurevich, L., de Jonge, N., and Nielsen, J.L. (2020). Impact of polyethylene on salivary glands proteome in Galleria melonella. Comp Biochem Physiol Part D Genomics Proteomics 34, 100678. 10.1016/j.cbd.2020.100678

34. Zhang, J., Gao, D., Li, Q., Zhao, Y., Li, L., Lin, H., Bi, Q., and Zhao, Y. (2020). Biodegradation of polyethylene microplastic particles by the fungus Aspergillus flavus from the guts of wax moth Galleria mellonella. Sci Total Environ 704, 135931. 10.1016/j.scitotenv.2019.135931

35. Cassone, B.J., Grove, H.C., Elebute, O., Villanueva, S.M.P., and LeMoine, C.M.R. (2020). Role of the intestinal microbiome in low-density polyethylene degradation by caterpillar larvae of the greater wax moth, Galleria mellonella. Proc Biol Sci 287, 20200112. 10.1098/rspb.2020.0112

36. Lou, Y., Ekaterina, P., Yang, S.S., Lu, B., Liu, B., Ren, N., Corvini, P.F., and Xing, D. (2020). Biodegradation of polyethylene and polystyrene by greater wax moth larvae (Galleria mellonella L.) and the effect of co-diet supplementation on the core gut microbiome. Environ Sci Technol 54, 2821–2831. 10.1021/acs.est.9b07044

37. LeMoine, C.M., Grove, H.C., Smith, C.M., and Cassone, B.J. (2020). A very hungry caterpillar: polyethylene metabolism and lipid homeostasis in larvae of the greater wax moth (Galleria mellonella). Environ Sci Technol 54, 14706–14715. 10.1021/acs.est.0c04386

38. Ren, L., Men, L., Zhang, Z., Guan, F., Tian, J., Wang, B., Wang, J., Zhang, Y., and Zhang, W. (2019). Biodegradation of polyethylene by Enterobacter sp. D1 from the guts of wax moth Galleria mellonella. Int J Environ Res Public Health 16. 10.3390/ijerph16111941

39. Dong, M., Zhang, Q., Xing, X., Chen, W., She, Z., and Luo, Z. (2020). Raman spectra and surface changes of microplastics weathered under natural environments. Sci Total Environ 739, 139990. 10.1016/j.scitotenv.2020.139990

40. Zhang, K., Hamidian, A.H., Tubic, A., Zhang, Y., Fang, J.K.H., Wu, C., and Lam, P.K.S. (2021). Understanding plastic degradation and microplastic formation in the environment: A review. Environ Pollut 274, 116554. 10.1016/j.envpol.2021.116554. 10.1016/j.envpol.2021.116554

41. Larkin, P. (2011) Infrared and Raman Spectroscopy; Principles and Spectral Interpretation. Elsevier: Waltham, MA.

42. Tromifchuk, E.S., Moskvina, M. A., Nikonorova, N. I., Efimov, A. V., Garina, E. S., Grokhovskaya, E. E., Ivanova, O. A., Bakirov, A. V. Sedush, N. G. and Chvalun, S. N. (2020). Hydrolytic degradation of polylactide films deformed by the environmental crazing mechanism. European Polymer Journal 139. 10.1016/j.eurpolymj.2020.110000

43. Cassone, B.J., Grove, H.C., Kurchaba, N., Geronimo, P., and LeMoine, C.M.R. (2022). Fat on plastic: Metabolic consequences of an LDPE diet in the fat body of the greater wax moth larvae (Galleria mellonella). J Hazard Mater 425, 127862. 10.1016/j.jhazmat.2021.127862

44. Montazer, Z., Habibi Najafi, M.B., and Levin, D.B. (2021). In vitro degradation of low-density polyethylene by new bacteria from larvae of the greater wax moth, Galleria mellonella. Can J Microbiol 67, 249–258. 10.1139/cjm-2020-0208

45. Réjasse, A.W. J., Deniset-Besseau, A., Crapart, N., Nielsen-Leroux, C. and Sandt, C. (2022). Plastic biodegradation: Do Galleria mellonella larvae bioassimilate polyethylene? A spectral histology approach using isotopic labeling and infrared microspectroscopy. Environmental Science and Technology 56, 525–534. 10.1021/acs.est.1c03417

46. Taghavi, N., Singhal, N., Zhuang, W.-Q. and Baroutian, S. (2020). Degradation of plastic waste using stimulated and naturally occurring microbial strains. Chemosphere 263. 10.1016/j.chemosphere.2020.127975

47. Zielin’ska, E., Zielin’ski, D., Jakubczyk, A., Kara’s, M., Pankiewicz, U., Flasz, B., Dziewięcka, M. and Lewicki, S. (2021). The impact of polystyrene consumption by edible insects Tenebrio molitor and Zophobas morio on their nutritional value, cytotoxicity, and oxidative stress parameters. Food Chemistry 345. 10.1016/j.foodchem.2020.128846

48. Tsochatzis, E., Lopes, J.A., Gika, H., and Theodoridis, G. (2020). Polystyrene biodegradation by Tenebrio molitor larvae: Identification of generated substances using a GC-MS untargeted screening method. Polymers (Basel) 13. 10.3390/polym13010017

49. Yang, S.S., Ding, M.Q., Zhang, Z.R., Ding, J., Bai, S.W., Cao, G.L., Zhao, L., Pang, J.W., Xing, D.F., Ren, N.Q., and Wu, W.M. (2021). Confirmation of biodegradation of low-density polyethylene in darkversus yellowmealworms (larvae of Tenebrio obscurus versus Tenebrio molitor) via. gut microbe-independent depolymerization. Sci Total Environ 789, 147915. 10.1016/j.scitotenv.2021.147915

50. Rojo, F. (2009). Degradation of alkanes by bacteria. Environ Microbiol 11, 2477–2490. 10.1111/j.1462-2920.2009.01948.x

51. Inderthal, H., Tai, S.L., and Harrison, S.T.L. (2021). Non-Hydrolyzable plastics -An interdisciplinary look at plastic bio-oxidation. Trends Biotechnol 39, 12–23. 10.1016/j.tibtech.2020.05.004

52. Hughes, A.L. (1999). Evolution of the arthropod prophenoloxidase/hexamerin protein family. Immunogenetics 49, 106–114. 10.1007/s002510050469

53. Glazer, L., Tom, M., Weil, S., Roth, Z., Khalaila, I., Mittelman, B., and Sagi, A. (2013). Hemocyanin with phenoloxidase activity in the chitin matrix of the crayfish gastrolith. J Exp Biol 216, 1898–1904. 10.1242/jeb.080945

54. Besser, K., Malyon, G.P., Eborall, W.S., Paro da Cunha, G., Filgueiras, J.G., Dowle, A., Cruz Garcia, L., Page, S.J., Dupree, R., Kern, M., et al. (2018). Hemocyanin facilitates lignocellulose digestion by wood-boring marine crustaceans. Nat Commun 9, 5125. 10.1038/s41467-018-07575-2

55. Rivera-Vega, L.J., Acevedo, F.E., and Felton, G.W. (2017). Genomics of Lepidoptera saliva reveals function in herbivory. Curr Opin Insect Sci 19, 61–69. 10.1016/j.cois.2017.01.002

56. Urbanska, A., Tjallingii, W. F., Dixon, A. F. G., Leszczynski, B. (1998). Phenol oxidising enzymes in the grain aphid’s saliva. Entomologia Experimentalis et Applicata 86, 197–203. 10.1046/j.1570-7458.1998.00281.x

57. Tian, D., Peiffer, M., Shoemaker, E., Tooker, J., Haubruge, E., Francis, F., Luthe, D.S., and Felton, G.W. (2012). Salivary glucose oxidase from caterpillars mediates the induction of rapid and delayed-induced defenses in the tomato plant. PLoS One 7, e36168. 10.1371/journal.pone.0036168

58. Bhattacharya, A., Sood, P., and Citovsky, V. (2010). The roles of plant phenolics in defence and communication during Agrobacterium and Rhizobium infection. Mol Plant Pathol 11, 705–719. 10.1111/j.1364-3703.2010.00625.x

59. Peng, L., Yan, Y., Yang, C. H., Barro, P. J. and Wan, F. (2013). Identification, comparison, and functional analysis of salivary phenol oxidizing enzymes in Bemisia tabaci B and Trialeurodes vaporariorum. Entomologia Experimentalis et Applicata 147, 282–292. 10.1111/eea.12068

60. Wu, K., Zhang, J., Zhang, Q., Zhu, S., Shao, Q., Clark, K.D., Liu, Y., and Ling, E. (2015). Plant phenolics are detoxified by prophenoloxidase in the insect gut. Sci Rep 5, 16823. 10.1038/srep16823

## References (Methods)

1. Anastassiades, M., Lehotay, S. J., Štajnbaher, D., and Schenck, F.J. (2019). Fast and easy multiresidue method employing acetonitrile extraction/partitioning and “dispersive solid-phase extraction” for the determination of pesticide residues in produce. Journal of AOAC INTERNATIONAL 86, 412–431. 10.1093/jaoac/86.2.412

2. Tsochatzis, E., Lopes, J.A., Gika, H., and Theodoridis, G. (2021). Polystyrene biodegradation by Tenebrio molitor larvae: identification of generated substances using a GC-MS untargeted screening method. Polymer 13. 10.3390/polym13010017

3. Kall, L., Canterbury, J.D., Weston, J., Noble, W.S., and MacCoss, M.J. (2007). Semi-supervised learning for peptide identification from shotgun proteomics datasets. Nat Methods 4, 923–925. 10.1038/nmeth1113

4. Dobin, A., Davis, C.A., Schlesinger, F., Drenkow, J., Zaleski, C., Jha, S., Batut, P., Chaisson, M., and Gingeras, T.R. (2013). STAR: ultrafast universal RNA-seq aligner. Bioinformatics 29, 15–21. 10.1093/bioin-formatics/bts635

5. Grabherr, M.G., Haas, B.J., Yassour, M., Levin, J.Z., Thompson, D.A., Amit, I., Adiconis, X., Fan, L., Raychowdhury, R., Zeng, Q., et al. (2011). Full-length transcriptome assembly from RNA-Seq data without a reference genome. Nature biotechnology 29, 644–652. 10.1038/nbt.1883

6. Brůna, T., Hoff, K.J., Lomsadze, A., Stanke, M., and Borodovsky, M. (2021). BRAKER2: automatic eukaryotic genome annotation with GeneMark-EP+ and AUGUSTUS supported by a protein database. NAR Genomics and Bioinformatics 3. 10.1093/nargab/lqaa108

7. Stanke, M., and Morgenstern, B. (2005). AUGUSTUS: a web server for gene prediction in eukaryotes that allows user-defined constraints. Nucleic acids research 33, W465–467. 10.1093/nar/gki458

8. Toronen, P., Medlar, A., and Holm, L. (2018). PANNZER2: a rapid functional annotation web server. Nucleic Acids Res 46, W84–W88. 10.1093/nar/gky350

9. Simão, F.A., Waterhouse, R.M., Ioannidis, P., Kriventseva, E.V., and Zdobnov, E.M. (2015). BUSCO: assessing genome assembly and annotation completeness with single-copy orthologs. Bioinformatics 31, 3210–3212. 10.1093/bioinformatics/btv351

